# Focal adhesion protein vinculin inhibits Mef2c-driven sclerostin expression in osteocytes to promote bone formation in mice

**DOI:** 10.1101/2022.11.09.515817

**Authors:** Yishu Wang, Jianmei Huang, Sixiong Lin, Lei Qin, Dingyu Hao, Peijun Zhang, Shaochuan Huo, Xuenong Zou, Di Chen, Guozhi Xiao

**Affiliations:** Department of Biochemistry, School of Medicine, Guangdong Provincial Key Laboratory of Cell Microenvironment and Disease Research, Shenzhen Key Laboratory of Cell Microenvironment, Southern University of Science and Technology, Shenzhen, 518055, China; Department of Spine Surgery, Orthopedic Research Institute, The First Affiliated Hospital of Sun Yat-sen University, Guangdong Provincial Key Laboratory of Orthopedics and Traumatology, Guangzhou 510080, China; Department of Orthopedics, Huazhong University of Science and Technology Union Shenzhen Hospital, Shenzhen 518000, China; Research Institute, Shenzhen Hospital (Futian) of Guangzhou University of Chinese Medicine, Shenzhen 518000, China; Research Center for Human Tissues and Organs Degeneration, Shenzhen Institutes of Advanced Technology, Chinese Academy of Sciences, Shenzhen 518055, China

**Keywords:** focal adhesion, vinculin, Mef2c, sclerostin, bone formation

## Abstract

The focal adhesion (FA) is the structural basis of the cell-extracellular matrix crosstalk and plays important roles in control of organ formation and function. Here we show that expression of FA protein vinculin is dramatically reduced in osteocytes in patients with aging-related osteoporosis. Deleting vinculin using the mouse 10-kb *Dmp1-Cre* transgenic mice causes a severe osteopenia in both young and aged mice by impairing osteoblast function without affecting bone resorption. Vinculin loss impairs the anabolic response of skeleton to mechanical loading in mice. At the molecular level, vinculin knockdown increases, while vinculin overexpression decreases, sclerostin expression in osteocytes without impacting expression of Mef2c, a major transcriptional regulator of the *Sost* gene, which encodes sclerostin. Vinculin interacts with Mef2c and vinculin loss enhances Mef2c nuclear translocation and binding to the *Sost* enhancer *ECR5* to promote sclerostin production in osteocytes. Deleting *Sost* expression reverses the osteopenic phenotypes caused by vinculin loss in mice. We demonstrate that estrogen is a major regulator of vinculin expression in osteocytes and that more importantly vinculin loss mediates estrogen deficiency induction of bone loss in mice. Thus, we demonstrate a novel mechanism through which vinculin inhibits the Mef2c-driven sclerostin expression in osteocytes to promote bone formation.

## Introduction

The adult skeleton is a dynamic tissue that undergoes constant bone remodeling, a process during which the osteoclast-mediated bone resorption is closely coupled with the osteoblast-mediated bone formation (Compston, McClung, & Leslie, 2019; X. Feng & McDonald, 2011). Impairment in the bone remodeling process results in osteoporosis characterized with low bone mass and bone fracture (Einhorn & Gerstenfeld, 2015; Wawrzyniak & Balawender, 2022). Understanding the mechanisms behind bone remodeling will advance bone biology research and identify new targets for treating metabolic bone diseases (Wong, Chin, Suhaimi, Ahmad, & Ima-Nirwana, 2016).

The focal adhesion (FA), which consists of integrins, extracellular matrix (ECM), cytoskeleton, kindlin-2, talin, vinculin and other proteins, mediates communication between cells and their environment and plays a central role in regulation of the cell-ECM adhesion, migration, differentiation, survival and mechanotransduction (Calderwood, Campbell, & Critchley, 2013; Larjava, Plow, & Wu, 2008; Ma, Qin, Wu, & Plow, 2008; Montanez et al., 2008; Qin, Liu, Cao, & Xiao, 2020; Qin, Yang, Yi, Cao, & Xiao, 2021; Ussar, Wang, Linder, Fässler, & Moser, 2006; H. Xu, Cao, & Xiao, 2016). Abnormalities in expression and activation of the FA proteins are involved in the pathogenesis and metastasis of cancers (Guo & Wu, 2019, 2020; Lin et al., 2017; Theodosiou et al., 2016; Xue, Xue, Wan, Li, & Shi, 2020; Zhan & Zhang, 2018). More recent studies reveal that FA proteins are critical for control of formation and organs and tissues, including skeleton (Cao et al., 2020; S. Chen et al., 2022; Fu et al., 2020; Lai et al., 2022; Lei et al., 2020; Qin, Fu, et al., 2021; Theodosiou et al., 2016; Y. Wang et al., 2019; C. Wu et al., 2015; X. Wu, Cao, & Xiao, 2021; X. Wu et al., 2022), heart (Liang, Sun, & Chen, 2009; Qi et al., 2019; Zhang et al., 2016), kidney (Lausecker et al., 2018; Sun et al., 2017; Wei et al., 2013), pancreatic islets (Zhu et al., 2020), adipose tissue (H. Gao et al., 2019), small intestine (He et al., 2020), testicle (Chi et al., 2021), and liver(H. Gao et al., 2021; H. Gao et al., 2022). Vinculin is a key FA protein that is highly concentrated when cells contact one another and the underlying substratum (Bays & DeMali, 2017; Burridge & Feramisco, 1980; Geiger, 1979). Vinculin is critical for the FA formation and maintenance. Cells overexpressing vinculin assemble large FAs (Janssen et al., 2006; Rodríguez Fernández, Geiger, Salomon, & Ben-Ze’ev, 1993), whereas vinculin-deficient cells form smaller and fewer FAs (Peng, Maiers, Choudhury, Craig, & DeMali, 2012; W. Xu, Coll, & Adamson, 1998). Vinculin exists in two conformations in the cell, i.e., an open, active form and a closed, auto-inhibited state in which the head domain forms extensive interactions with the tail (H. Chen, Choudhury, & Craig, 2006; Izard et al., 2004). Mechanical stimulation alters the morphology of vinculin. Vinculin also responds to force and conducts to the cytoskeleton (Golji, Wendorff, & Mofrad, 2012; Grashoff et al., 2010). Previous studies have demonstrated that vinculin is necessary for the development and homeostasis of platelets (Mitsios et al., 2010), neocortical neurons (Mandal, Belapurkar, Nair, & Ramanan, 2021), kidney (Lausecker et al., 2018; Palovuori & Eskelinen, 2000), pancreatic islets (Shi, Guo, Li, Wang, & Qin, 2018; Y. Wang et al., 2012), stomach (Kim et al., 2020; Li et al., 2021) and breast (Y. Gao et al., 2017). However, the function of vinculin in bone has not been explored.

In this study, we demonstrate that mice lacking vinculin in the dentin matrix protein 1 (Dmp1)-positive cells (i.e., osteocytes and mature osteoblasts) display a severe bone loss with impaired bone formation, fail to properly respond to mechanical loading, and do not show further bone loss on estrogen deficiency. Vinculin interacts with Mef2c and vinculin loss increases Mef2c nuclear translocation and binding to the *Sost* enhancer *ECR5* to promote sclerostin production in osteocytes and thereby inhibit bone formation.

## Results

### Vinculin is downregulated in osteocytes in human osteoporotic bones and vinculin loss impairs osteocyte adhesion and spreading

As an initial attempt to explore the potential involvement of alteration in expression of FA protein vinculin in the pathogenesis of osteoporosis, we collected human bone tissue samples from young (from 29 to 32 years old) and aged individuals (from 78 to 88 years old) and performed immunofluorescence (IF) staining of these bone sections to determine the expression level of vinculin protein. The results showed that osteocytes embedded in the trabecular bone matrix in the young bone samples expressed a high level of vinculin protein, which was drastically decreased in the aged bones (Fig. 1a, b). To gain insights into the function of vinculin in osteocytes, we utilized the CRISPR-Cas9 technology to delete the *Vcl* gene, which encodes vinculin, in MLO-Y4 osteocyte-like cells. We found that MLO-Y4 cells with deletion of two alleles of the *Vcl* gene did not grow and die in culture (data not shown). We therefore knocked down (KD) vinculin expression by deleting one allele of the *Vcl* gene in MLO-Y4 cells. Note: vinculin KD did not markedly affect the expression levels of other FA proteins, such as integrinβ1, kindlin-2, talin and pinch1 (Fig. 1c, d), as measured by IF staining and western blotting analyses. Vinculin KD greatly altered the distribution and orientation of filamentous actin (Fig. 1e) and impaired cell spreading in MLO-Y4 cells (Fig. 1f).

**Figure 1.**
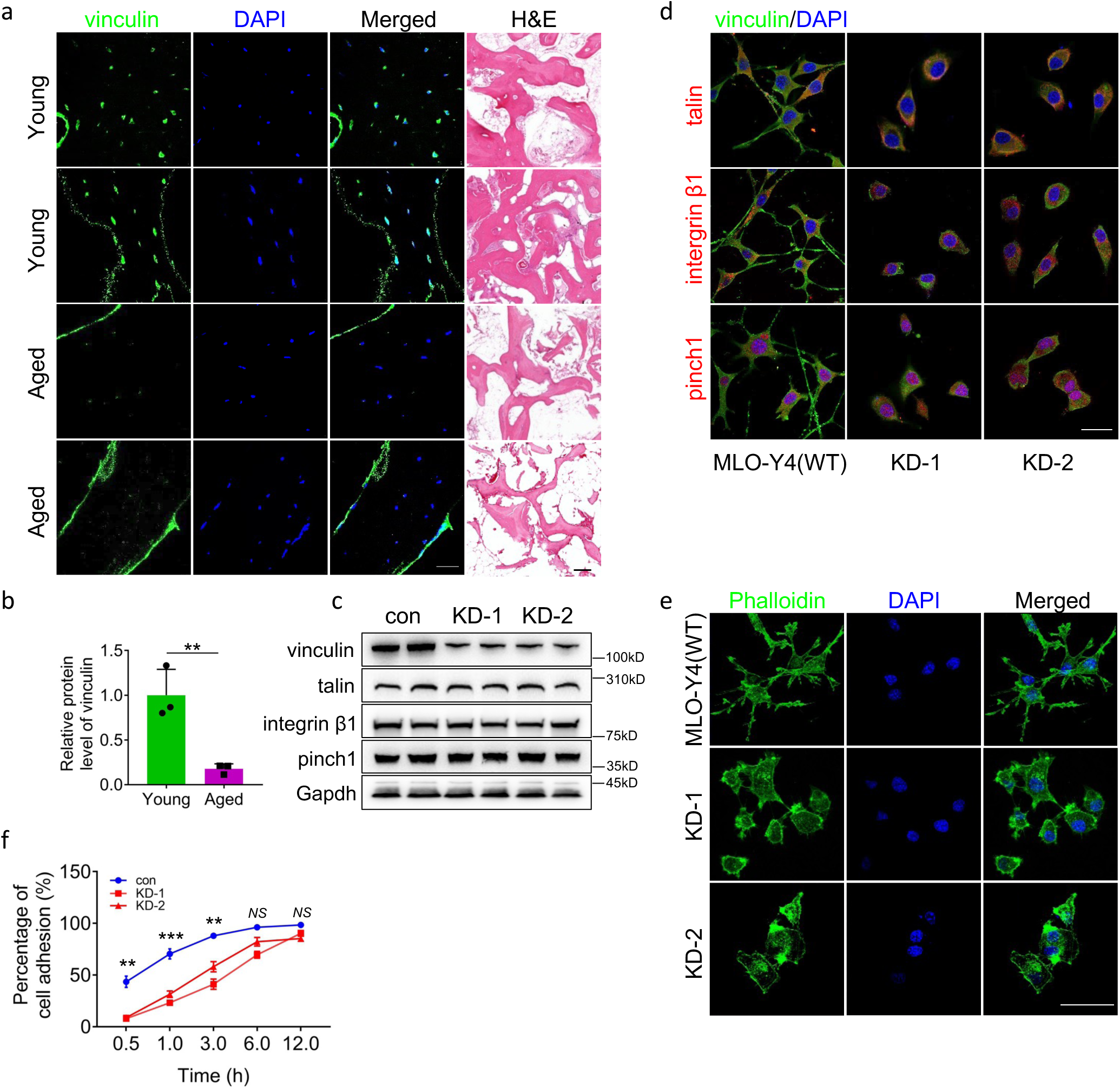
Vinculin regulates osteocyte adhesion and is downregulated in osteocytes in human osteoporotic bones. **a, b** Immunofluorescence (IF) staining and hematoxylin and eosin (H/E) staining. Sections of human cancellous samples from young (29-32-yrs-old) and old 78-88-yrs-old) subjects were subjected to H/E and IF staining with a vinculin antibody. Scale bars: 50 μm. Quantitative data (b). Results were expressed as mean ± s.d. *N* = 3 biologically independent replicates per group, *\*\*P* < 0.01 versus young, unpaired two-tailed Student’s *t* test. **c** Western blot analysis. Protein extracts isolated from MLO-Y4 osteocyte-like cells with and without vinculin knockdown (KD) by CRISPR-Cas9 technology were subjected to western blotting using the indicated antibodies. **d** IF staining. MLO-Y4 cells with and without vinculin KD were subjected to IF staining using phalloidin or DAPI. **e** IF staining. MLO-Y4 cells with and without vinculin KD were subjected to IF staining using the indicated antibodies. **f** Cell adhesion assay. MLO-Y4 cells with and without vinculin KD were seeded in a 96-well plate at a density of 1×10^5^ cells/well. The absorbance was measured at the time points of 0.5h, 1h, 3h, 6h and 12h, respectively. Results were expressed as mean ± s.d., \*\**P* < 0.01, \**\*\*P* < 0.001 versus controls, unpaired two-tailed Student’s *t* test.

### Deleting vinculin expression in Dmp1-expressing cells causes a low bone mass in mice

To investigate potential role of vinculin in bone, we deleted its expression in the dentin matrix protein 1 (Dmp1)-expressing cells (primarily osteocytes and mature osteoblasts) by breeding the floxed *Vcl* mice (*Vcl^fl/fl^*) with the 10-kb mouse *Dmp1-Cre* transgenic mice (*Dmp1-Cre; Vcl^fl/fl^*, referred as cKO hereafter). The results from qRT-PCR and western blotting analyses showed that vinculin mRNA and protein expression was significantly reduced in cKO cortical bones compared to that in control bones (Fig. 2a, b). Micro-computed tomography (μCT) analysis of distal femurs from 3- and 14-mo-old male control and cKO mice showed that vinculin loss caused a severe osteopenia at both ages. Vinculin loss significantly decreased the bone mineral density (BMD) (Fig. 2c, d) and bone volume fraction (BV/TV) (Fig. 2c, e) without affecting the cortical thickness (Ct.Th) (Fig. 2f). Consistent with results from μCT analyses, results from H/E staining of tibial sections showed much less trabecular bone mass in cKO mice than that in control mice (Fig. 2g). Similar reductions in BMD (Fig. 2h, i) and BV/TV (Fig. 2h, j) were observed in 3-mo-old male cKO spine (L4-5) compared to those in control mice (Fig. 2h-j). Note: vinculin loss did not affect the bone mass in skull (Fig. 2k).

**Figure 2.**
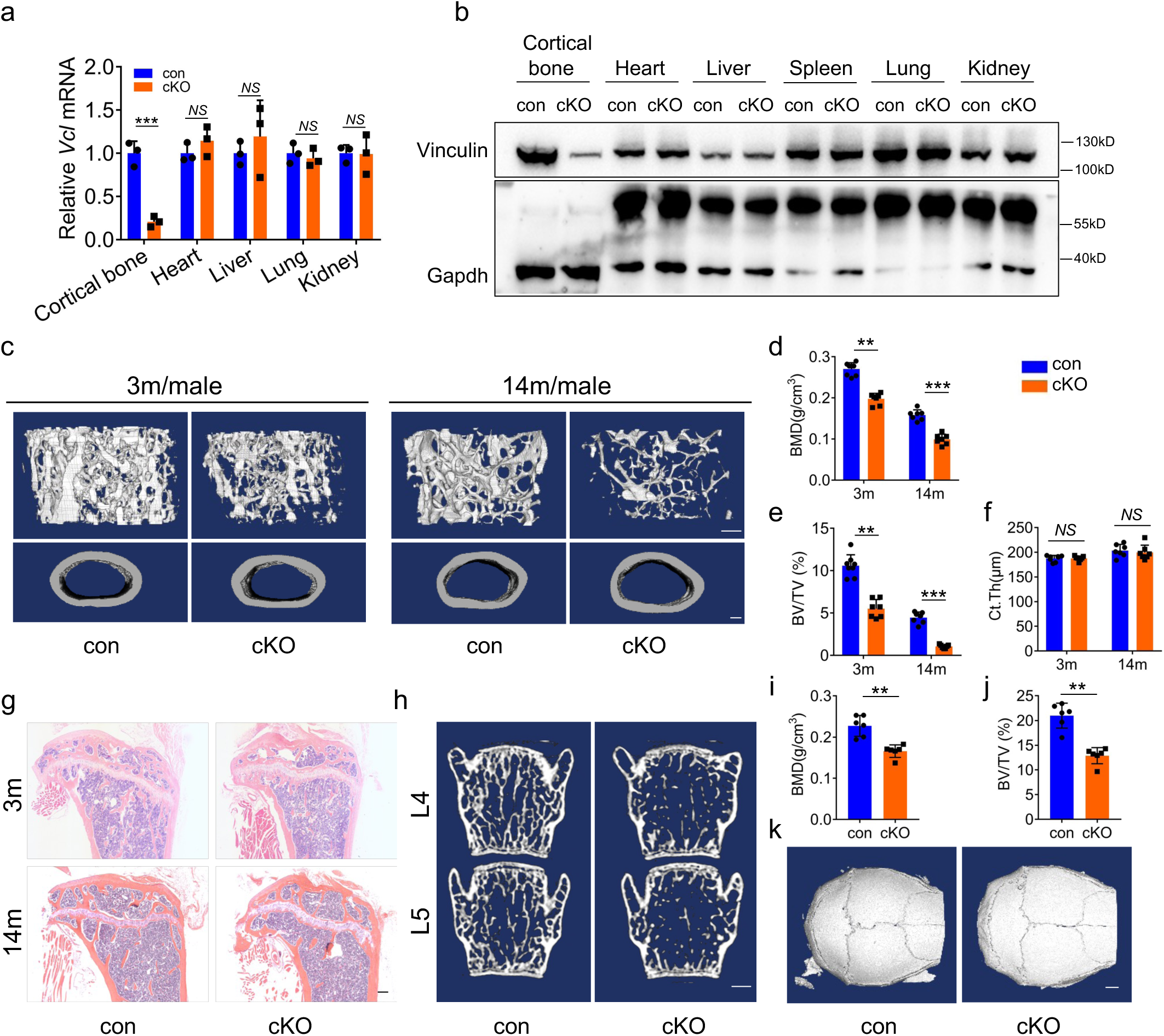
Vinculin loss in Dmp1-expressing cells causes severe osteopenia by impairing bone formation in mice. **a** Real-time RT-PCR (qPCR) analyses. Total RNAs isolated from the indicated tissues of 3 mo-old male control (con) and cKO mice were used for qPCR analysis for expression of *Vcl* gene, which was normalized to *Gapdh* mRNA. Results were expressed as mean ± s.d., *N* = 3 biologically independent replicates per group, *\*\*P* < 0.01 versus controls, unpaired two-tailed Student’s *t* test. **b** Western blot analysis. Protein extracts were isolated from the indicated tissues of the control and cKO mice were subjected to western blotting analysis for vinculin expression. Gapdh was used for loading control. **c** Three-dimensional (3D) reconstruction from micro-computerized tomography (μCT) scans of distal femurs from male control and cKO mice with the indicated ages. **d-f** Quantitative analyses of the bone volume/tissue volume (BV/TV), cortical thickness (Ct.Th) and bone mineral density (BMD) of distal femurs from male control and cKO mice with the indicated ages. Results were expressed as mean ± s.d., *N* = 6 biologically independent replicates per group, \**P* < 0.05, \**\*P* < 0.01, *\*\*\*P* < 0.001 versus controls, unpaired two-tailed Student’s *t* test. **g** H/E staining of tibial sections of 3-mo-old male control and cKO mice with the indicated ages. **h** 3D reconstruction from μCT scans of spine (L4, L5) from 3-mo-old male control and cKO mice. **i, j** Quantitative analyses of the bone volume/tissue volume (BV/TV) and bone mineral density (BMD) of spine from male control and cKO mice. Results were expressed as mean ± s.d., *N* = 6 biologically independent replicates per group, \**P* < 0.05, \**\*P* < 0.01 versus controls, unpaired two-tailed Student’s *t* test. **k** 3D reconstruction from μCT scans of skull.

### Vinculin loss mainly impairs bone formation without markedly impacting bone resorption

To investigate mechanism through which vinculin loss causes a low bone mass, we determined the osteoblast and osteoclast formation and function in control and cKO long bones. Results from the double calcein labeling experiments showed that the mineralization apposition rate (MAR) and bone formation rate (BFR) in the femur metaphyseal cancellous bones from 3-mo-old male cKO mice were significantly decreased compared to those in control mice (Fig. 3a-c). The serum level of the procollagen type 1 amino-terminal propeptide (P1NP), a marker of in vivo bone formation(D’Oronzo, Brown, & Coleman, 2017), was significantly lower in cKO mice than that in control mice (Fig. 3d). To determine the effects of vinculin loss on the osteoid production and mineralization, we performed the von Kossa staining of femur sections of control and dKO mice and found that the osteoid volume/tissue volume (OV/TV) and mineralized bone volume/tissue volume (mBV/TV) were decreased in cKO mice relative to those in control mice (Fig. 3e-g). We next performed the colony forming unit-fibroblast (CFU-F) assay and colony forming unit-osteoblast (CFU-OB) assay. The results showed that the number of CFU-OB, but not that of CFU-F, was reduced in cKO group compared to that in control group (Fig. 3h, i). We further measured the osteoclast formation and bone-resorbing activity in control and cKO mice. The serum levels of collagen type I cross-linked C-telopeptide (CTX), degradation products from type I collagen during in vivo osteoclastic bone resorption(D’Oronzo et al., 2017), were not significantly different between the two genotypes (Fig. 3j). Results from TRAP staining of tibial sections showed that the osteoclast surface/bone surface (Oc.S/BS) and osteoclast number/bone perimeter (Oc.Nb/BPm) in both primary and secondary cancellous bones were not markedly altered in cKO mice relative those in control mice (Fig. 3k-o).

**Figure 3.**
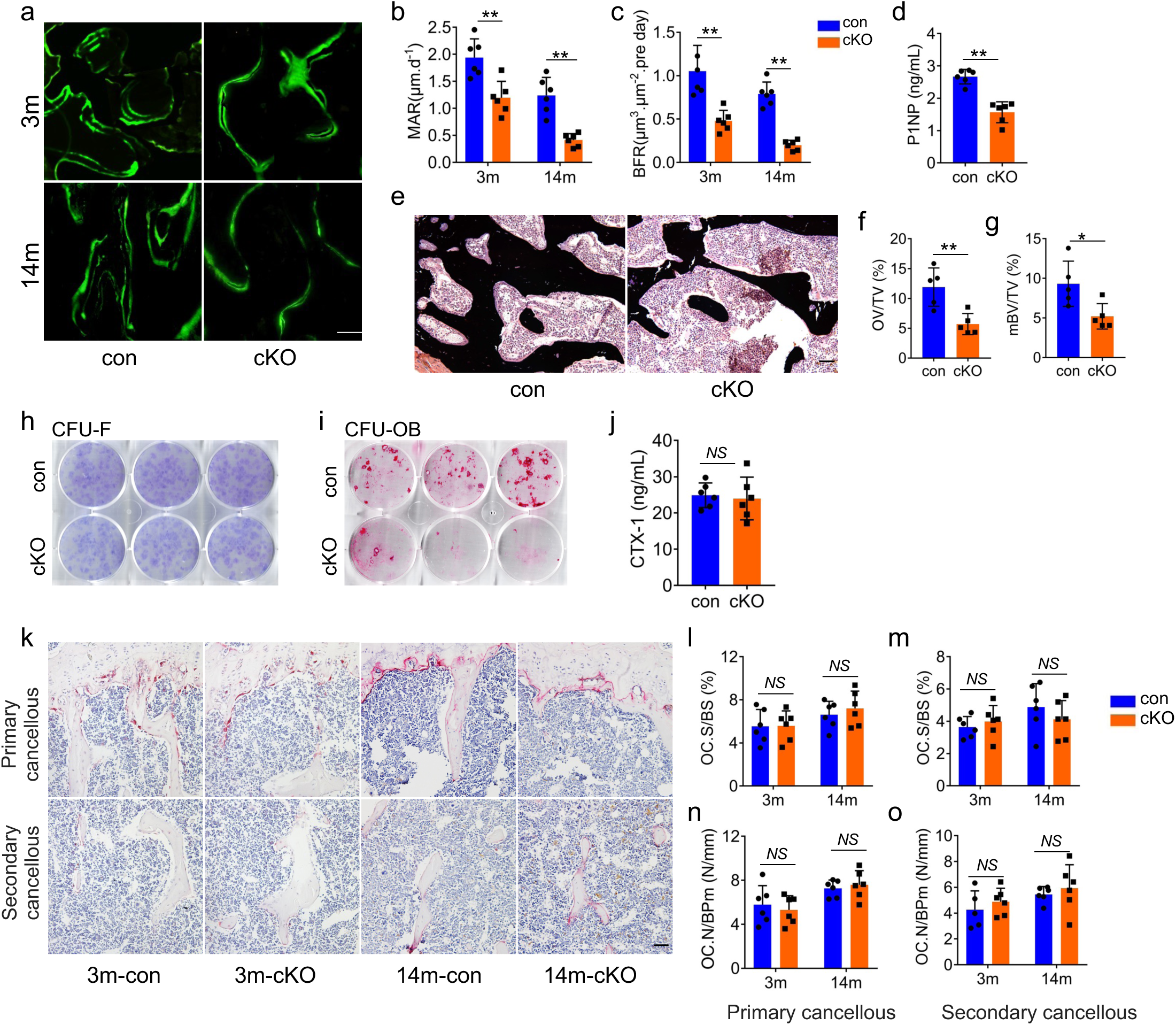
Vinculin loss impairs the osteoblast-mediated bone formation without affecting the osteoclast formation and bone resorption. **a-c** Calcein double labeling. Representative images of 3-mo-old male control and cKO femur sections (a). Sections of non-demineralized femurs of 3-mo-old male control and cKO mice were used for measurements of the mineral apposition rate (MAR) and bone formation rate (BFR). Quantitative MAR (b) and BFR (c) for the metaphyseal trabecular bones of the two genotypes. Scale bar, 50 μm. Results were expressed as mean ± s.d., *N* = 6 biologically independent replicates per group, \**P* < 0.05, \**\*P* < 0.01 versus controls, unpaired two-tailed Student’s *t* test. **d** Serum level of procollagen type 1 amino-terminal propeptide (P1NP). Sera harvested from 3-mo-old male control and cKO mice were subjected to ELISA assay for P1NP. Results were expressed as mean ± s.d., *N* = 6 biologically independent replicates per gro*up*, \**\*P* < 0.01 versus controls, unpaired two-tailed Student’s *t* test. **e-g** von Kossa staining. Undecalcified sections of femora from 3-mo-old male control and cKO mice were subjected to von Kossa staining (e). Quantitative OV/TV (osteoid volume/tissue volume) (f) and mBV/TV (mineralized bone volume/tissue volume) (g) data for the cancellous bones from distal femora were measured by bone morphometry. Scale bar, 50 μm. Results were expressed as mean ± s.d., *N* = 5 biologically independent replicates per group, \**P* < 0.05, \**\*P* < 0.01 versus controls, unpaired two-tailed Student’s *t* test. **h** Colony forming unit-fibroblast (CFU-F) assays. Bone marrow nucleated cells from 3-mo-old male control and cKO mice were seeded in a 6-well plate with a cell density of 2×10^6^ per well and cultured using the Mouse MesenCult proliferation kit (CFU-F assay) for 14 days, followed by Giemsa staining. **i** Colony forming unit-osteoblast (CFU-OB) assays. Bone marrow nucleated cells from 3-mo-old male control and cKO mice were seeded in a 6-well plate with a cell density of 4×10^6^ per well and cultured in osteoblast differentiation medium (complete α-MEM containing 50 μg/ml L-ascorbic acid and 2.0 mM β-glycerophosphate) for 21 d and media were changed every 48h, followed by alizarin red staining. **j** Serum level of collagen type I cross-linked C-telopeptide (CTX). Sera collected from 3-mo-old male control and cKO mice were subjected to ELISA assay for CTX. Results were expressed as mean ± s.d., *N* = 6 biologically independent replicates per group, *\*P* < 0.05 versus controls, unpaired two-tailed *t* test. **k-o** Tartrate-resistant acid phosphatase (TRAP) staining. Tibial sections of 3-mo-male control and cKO mice were used for TRAP staining (k). Osteoclast surface/bone surface (Oc.S/BS) (l, m) and osteoclast number/bone perimeter (Oc.N/BPm) (n, o) of primary and secondary cancellous bones were measured using Image-Pro Plus 7.0. Scale bar, 50 μm. Results were expressed as mean ± s.d., *N* = 7 biologically independent replicates per group, *\*P* < 0.05 versus controls, unpaired two-tailed Student’s *t* test.

### Vinculin loss increases, while vinculin overexpression decreases, sclerostin expression without affecting the level of Mef2c protein in osteocytes

Sclerostin is encoded by *Sost* and mainly secreted by osteocytes. Sclerostin decreases osteoblast and bone formation via inhibiting the Wnt/μ-catenin signaling. Sclerostin loss or pharmacological neutralization greatly increases bone formation and bone mass (Reid & Billington, 2022). Because the osteoblast and bone formation was severely impaired by vinculin loss in the cKO mice, as demonstrated above, we next determined whether vinculin loss increased sclerostin production in mice. Results from by IHC, RT-qPCR and western blotting analyses revealed that the expression levels of sclerostin mRNA and protein were strikingly increased in cKO versus control osteocytes and bones (Fig. 4a-e). It should be noted that vinculin loss did not markedly alter the level of Mef2c protein in bone extracts (Fig. 4d, e), a major transcriptional regulator of the *Sost* gene. In contrast, overexpression of vinculin decreased the level of sclerostin in MLO-Y4 cells (Fig. 4f, g).

**Figure 4.**
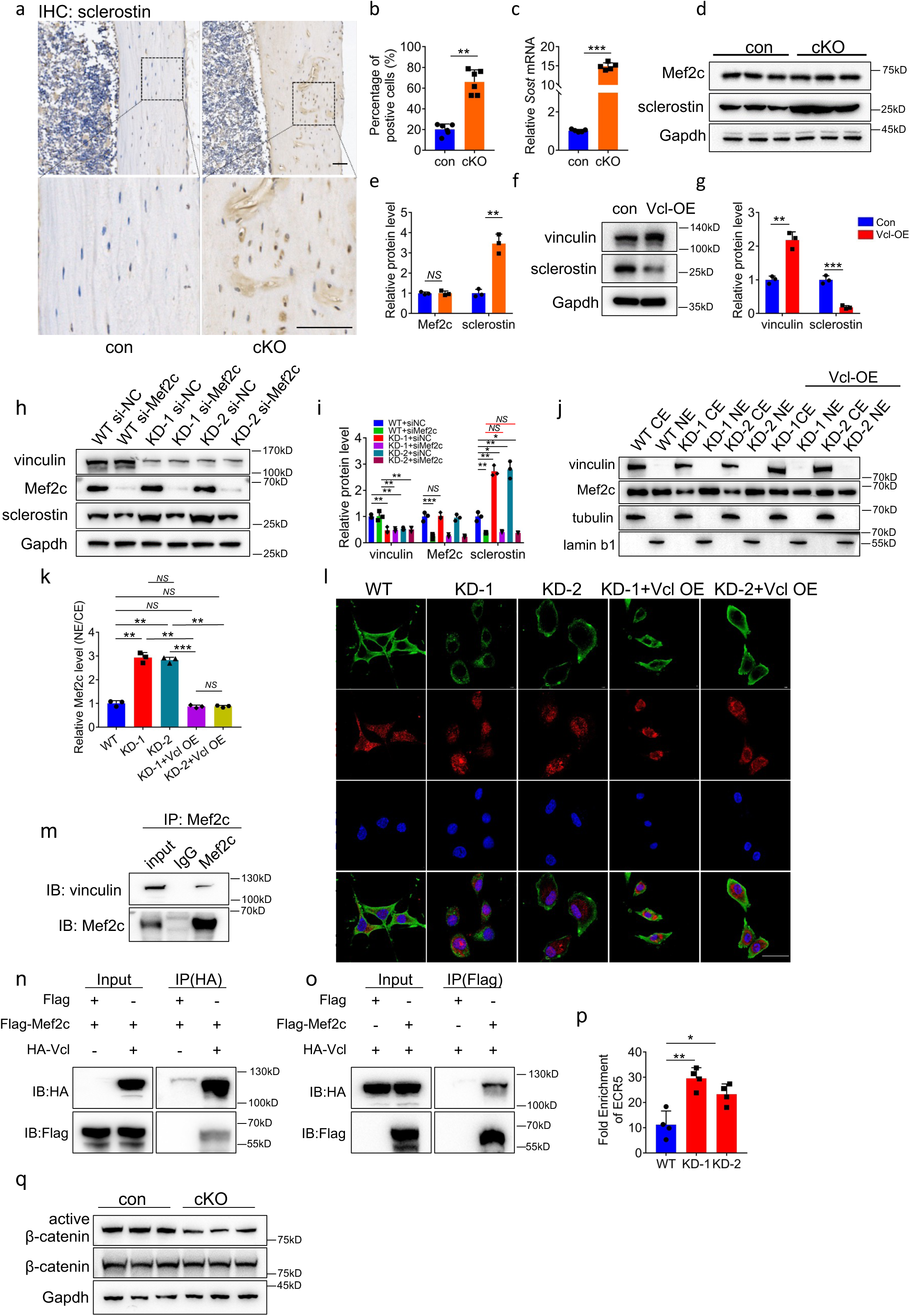
Vinculin interacts with Mef2c and vinculin knockdown increases Mef2c nuclear translocation and binding to the *Sost* enhancer *ECR5* to promote sclerostin expression in osteocytes. **a, b** Immunohistochemical (IHC) staining. Tibial sections of 3-mo-old male control and cKO mice were stained with an anti-sclerostin antibody. Quantitative data (b). *N* = 5 biologically independent replicates per group, *\*\*P* < 0.01, versus controls, unpaired two-tailed Student’s *t* test. **c** qPCR analyses. Total RNA isolated from cortical bone of 3 mo-old male mice was used for qPCR analysis for expression of *Sost* gene, which was normalized to *Gapdh* mRNA. Results were expressed as mean ± s.d., *N* = 5 biologically independent replicates per group, *\*\*\*P* < 0.001 versus controls, unpaired two-tailed Student’s *t* test. **d, e** Western blot analyses. Protein extracts isolated from cortical bones of 3 mo-old male mice were subjected to western blotting using the indicated antibodies. Results were expressed as mean ± s.d., *N =* 3 biologically independent replicates per group, \**\*P <* 0.01 versus controls, unpaired two-tailed Student’s *t* test. **f, g** Vinculin overexpression. Protein extracts isolated from MLO-Y4 cells transfected with control or vinculin-expressing plasmid were subjected to western blotting using the indicated antibodies. Results were expressed as mean ± s.d., \**\*P* < 0.01, \**\*\*P* < 0.001 versus veh, unpaired two-tailed Student’s *t* test. **h, i** Whole cell extracts from MLO-Y4 cells with and without vinculin or Mef2c siRNA knockdown (KD) were subjected to western blotting using the indicated antibodies. Results were expressed as mean ± s.d., *\*P* < 0.05, \**\*P* < 0.01 versus veh, two-way ANOVA**. j,k** Mef2c nuclear translocation. Cytoplasmic (CE) and nuclear extracts (NE) from MLO-Y4 cells with and without vinculin KD and with and without vinculin overexpression (OE) were subjected to western blotting with the indicated antibody. Quantitative data (k). Results were expressed as mean ± s.d., *N* = 3 biologically independent replicates per group, *\*\*P* < 0.01, *\*\*\*P* < 0.001 versus controls, unpaired two-tailed Student’s *t* test. **l** Vinculin-Mef2c colocalization. MLO-Y4 cells treated as in h were subjected to IF staining with the indicated antibody. Scale bars: 50 μm**. m-o** Co-immunoprecipitation (co-IP) assay. Protein extracts from MLO-Y4 cells with and without overexpression of HA-vinculin and Flag-Mef2c were incubated with the indicated antibodies or IgG, and the immunocomplexes were separated by SDS-PAGE, followed by western blotting with the indicated antibodies. **p** CHIP-qPCR analysis. The expression of *Sost* enhancer *ECR5* combined with Mef2c antibody, which was normalized to IgG. Results were expressed as mean ± s.d., *\*P* < 0.05, \**\*P* < 0.01 versus control, unpaired two-tailed Student’s *t* test. **q** Western blot analyses. Protein extracts isolated from BMSC cultures of 3 mo-old male control and cKO mice were subjected to western blotting using the indicated antibodies.

### Vinculin interacts with Mef2c and vinculin loss enhances Mef2c nuclear translocation and binding to the *Sost* enhancer *ECR5* in osteocytes

We found that siRNA knockdown (KD) of Mef2c in MLO-Y4 cells reversed the sclerostin upregulation caused by vinculin loss (Fig. 4h, i). Furthermore, vinculin KD decreased the level of cytoplasmic Mef2c and, in the meantime, increased that of its nuclear protein in MLO-Y4 cells, which were reversed by vinculin overexpression (Fig. 4j, k). IF staining revealed that vinculin and Mef2c colocalized in MLO-Y4 cells and that vinculin KD increased Mef2c nuclear translocation in these cells, which was reversed by vinculin overexpression (Fig. 4l). Results from co-immunoprecipitation (Co-IP) experiments demonstrated interactions of endogenous or overexpressed vinculin and Mef2c proteins in MLO-Y4 cells (Fig. 4m-o). Results from ChIP assay using the MLO-Y4 cells revealed that vinculin KD dramatically increased the binding of Mef2c to the *Sost* enhancer *ECR5* (Fig. 4p). Consistent with the fact that sclerostin is a potent inhibitor of the Wnt/μ-catenin pathway, the level of active μ-catenin, but not that of its total protein, was dramatically reduced in cKO BMSC cultures relative to that in control BMSC cultures (Fig. 4q).

### Deleting *Sost* expression in Dmp1-expressing cells reverses the osteopenia induced by vinculin loss

Above results suggest that the sclerostin upregulation by vinculin loss may play a role in causing the bone loss in cKO mice. To determine if this is the case, we next determined the effects of *Sost* deletion in Dmp1-expressing cells on the osteopenic phenotypes caused by vinculin loss in mice. We bred the *Dmp1-Cre; Vcl^fl/fl^* (cKO) mice with *Sost^fl/fl^*mice and generated *Dmp1-Cre; Vcl^fl/fl^; Sost^fl/fl^* and *Dmp1-Cre; Sost^fl/fl^* mice. We performed μCT analyses of the femurs of the indicated groups (Fig. 5a). As expected, *Sost* loss increased the BMD, BV/TV and Ct.Th (Fig. 5b-d). Importantly, *Sost* deletion largely reversed the osteopenia of cKO mice (Fig. 5b, c). H/E staining of the tibial sections revealed that *Sost* ablation reversed the low bone mass in vinculin-deficient mice (Fig. 5e). *Sost* inactivation similarly reversed the osteopenic phenotypes in spine (L4-L5) in vinculin cKO mice (Fig. 5f-h). Because it is known that sclerostin reduces bone formation by inhibiting Wnt/μ-catenin pathway, we measured the bone-forming activity of osteoblasts in vivo by performing the double calcein labeling experiments. As expected, we observed significant increases in the MAR and BFR after *Sost* inactivation (Fig. 5i-k). Importantly, deleting *Sost* expression largely restored the impairment in bone formation in cKO mice (Fig. 5i-k). Results from the TRAP staining of tibial sections showed no marked differences in osteoclast formation between the control and cKO mice with and without *Sost* deletion (Fig. 5l-p).

**Figure 5.**
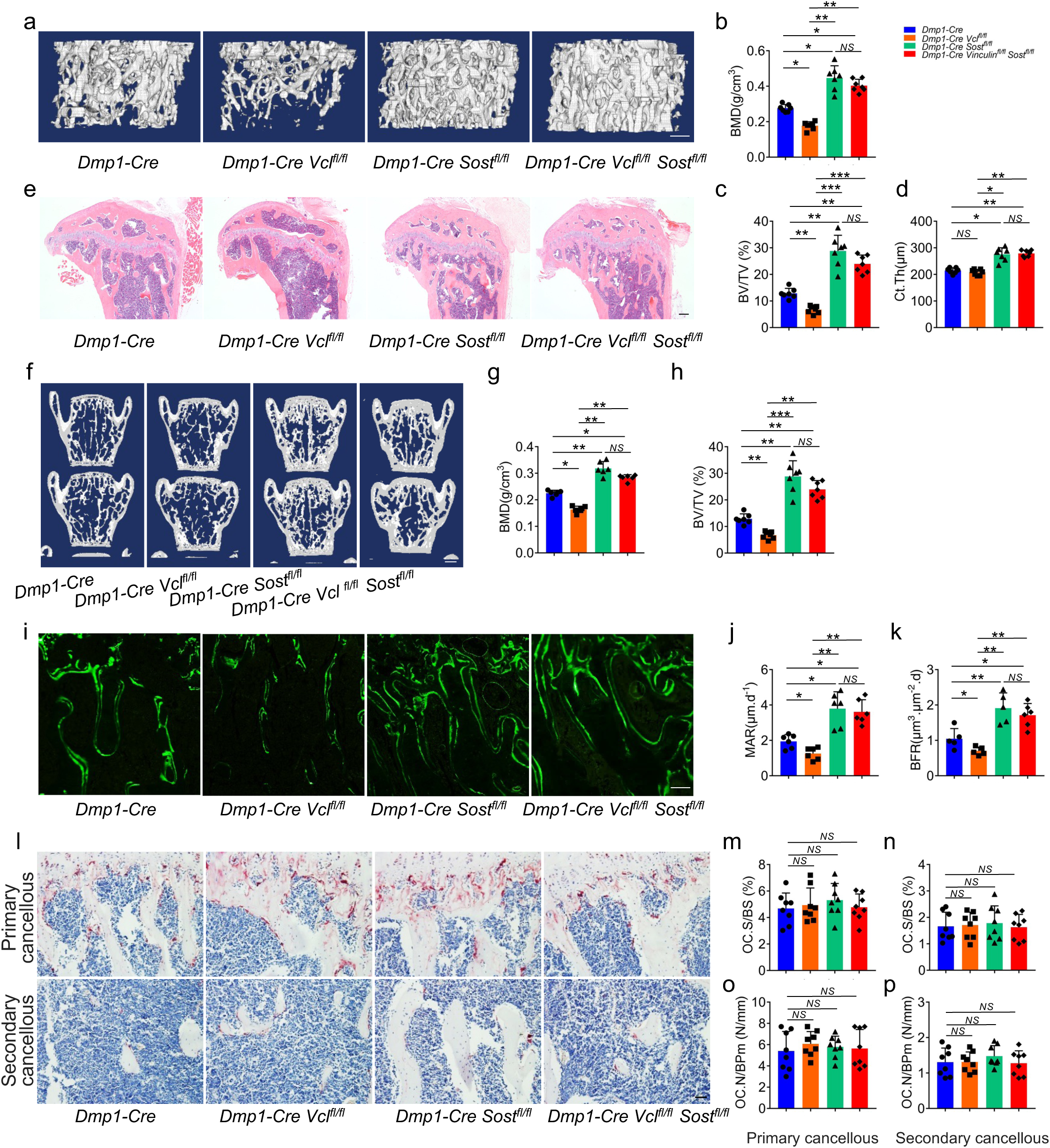
Deleting sclerostin in Dmp1-expressing cells reverses the osteopenia induced by vinculin loss. **a** 3D reconstruction from µCT scans of tibia from 3-mo-old male control and cKO mice treated with or without tibial loading experiment. Scale bar, 250 μm. **b** H/E staining of tibial sections of 3-mo-old control and cKO mice were treated with or without tibial loading experiment. Scale bar, 200 μm. **c-e** Quantitative analyses of BV/TV Ct.Th and BMD. Results were expressed as mean ± s.d., *N* = 7 biologically independent replicates per group, *\*P* < 0.05, \**\*P* < 0.01, \*\**\*P* < 0.001 versus controls, unpaired two-tailed Student’s *t* test. **f-h** Calcein double labeling. Representative images of femur sections (f). Sections were used for measurements of mineral apposition rate (MAR). Quantitative MAR data for metaphyseal trabecular bones (g) and bone formation rate (BFR) (h). Scale bar, 50 μm. **(i-m)** Tartrate-resistant acid phosphatase (TRAP) staining. Tibial sections of 3-mo-male mice were used for TRAP staining with indicated genotype. (i). Osteoclast surface/bone surface (Oc.S/BS) (j, k) and osteoclast number/bone perimeter (Oc.N/BPm) (l, m) of primary and secondary cancellous bones were measured using Image-Pro Plus 7.0. Scale bar, 50 μm. Results were expressed as mean ± s.d., *N* = 6 biologically independent replicates per group, *\*P* < 0.05 versus controls, unpaired two-tailed Student’s *t* test. **p** 3D reconstruction from μCT scans of spine (L4, L5) from 3-mo-old male mice with indicated genotype. **q, r** Quantitative analyses of bone volume/tissue volume (BV/TV) and bone mineral density (BMD) of spine. Results were expressed as mean ± s.d., *N* = 6 biologically independent replicates per group, \**P* < 0.05, \**\*P* < 0.01, \*\**\*P* < 0.001 versus controls, unpaired two-tailed Student’s *t* test.

### Vinculin loss impairs the anabolic responses of skeleton to mechanical loading in mice

It is now widely believed that osteocytes embedded in the mineralizing matrix are the major mechanosensors of bone (Lewis, 2021; Qin et al., 2020). We next examined whether vinculin is involved in the bone mechanotransduction by performing the tibial loading experiments in control and cKO mice (Qin et al., 2022). Results from μCT analyses showed that 2 weeks’ mechanical loading caused significant increases in the BMD, BV/TV and Ct.Th in control tibiae, which were essentially abolished in cKO tibiae (Fig. 6a-d). Results from H/E staining of the tibial sections showed that mechanical loading markedly increased the trabecular volume in control but not cKO mice (Fig. 6e). Mechanical loading significantly increased the values of MAR and BFR in control but not cKO tibiae (Fig. 6f-h). Finally, tibial loading decreased the Oc.Nb/BPm, but not the OC.N/BPm, in control mice, which was lost in cKO mice (Fig. 6i-m).

**Figure 6.**
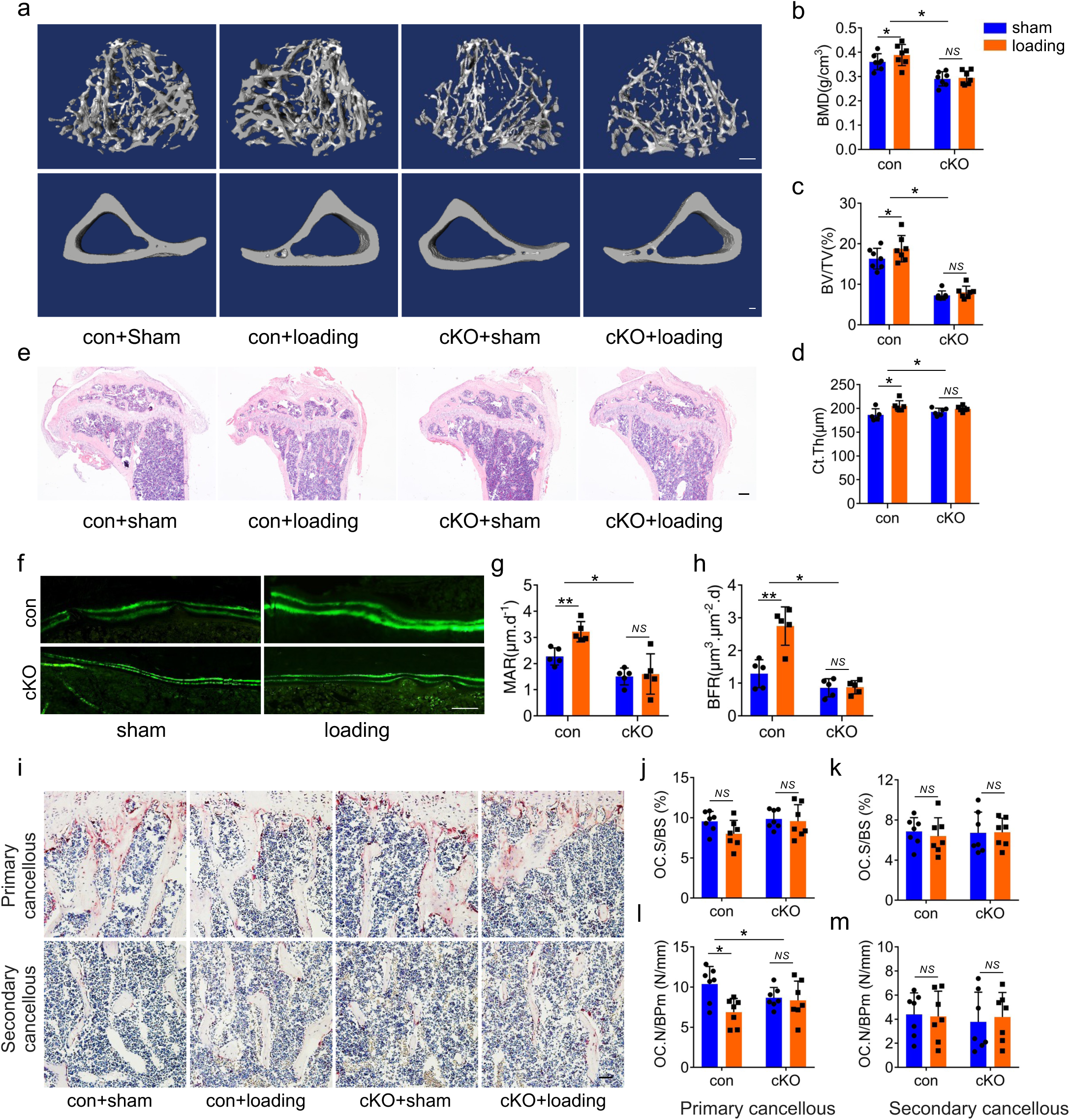
Vinculin deletion reduces mechanical loading-stimulated bone formation in mice. **a** 3D reconstruction from μCT scans of tibiae from 3-mo-old male control and cKO mice treated with or without tibia loading experiment. Scale bar, 100 μm. **b** H/E staining of tibial sections of 4-mo-old control and cKO mice were treated with or without tibia loading experiment. Scale bar, 200 μm. **c-e** Quantitative analyses of BV/TV, Ct.Th and BMD. Results were expressed as mean ± s.d., *N* = 7 biologically independent replicates per group, *\*P* < 0.05, \**\*P* < 0.01 versus controls, two-way ANOVA. **f-h** Calcein double labeling. Representative images of tibial sections (f). Sections were used for measurements of MAR and BFR for metaphyseal trabecular bones. Scale bar, 50 μm. Results were expressed as mean ± s.d., *N* = 5 biologically independent replicates per group, *\*P* < 0.05, \**\*P* < 0.01 versus controls, two-way ANOVA. **i-m** TRAP staining. Tibial sections of 4-mo-male control and cKO mice were used for TRAP staining (i). Oc.S/BS) (j, k) and Oc.N/BPm (l, m) of primary and secondary cancellous bones were measured using Image-Pro Plus 7.0. Scale bar, 50 μm. Results were expressed as mean ± s.d., *N* = 6 biologically independent replicates per group, *\*P* < 0.05 versus controls, two-way ANOVA.

### Estrogen loss reduces vinculin expression in osteocytes and ovariectomy no longer causes bone loss in vinculin-deficient mice

We next determined vinculin expression in osteocytes in mice with ovariectomy (OVX), which mimics the postmenopausal osteoporosis in humans. Surprisingly, we found that the expression level of vinculin protein was strikingly decreased in osteocytes embedded in the bone matrix in OVX mice (Fig. 7a, b). We next determined whether estrogen can directly regulate vinculin expression in cultured osteocytes and found that estrogen increased the level of vinculin protein in MLO-Y4 cells in a dose-dependent manner (Fig. 7c, d). In contrast, fulvestrant, an estrogen receptor antagonist, decreased the level of vinculin protein in MLO-Y4 cells (Fig. 7c, e). We next compared the effects of OVX on reduction of bone mass in control and vinculin cKO mice. To this end, 3-month-old control and vinculin cKO female mice were subjected to sham or OVX surgery, as we previously described (Fu et al., 2020). At two months after surgery, the bone mass was reduced in control mice, which is expected. Shockingly, the OVX-induced trabecular bone loss was dramatically reduced (BV/TV) or completely abolished (BMD) in cKO mice (Fig. 7f-i). While OVX did not markedly impact the MAR and BFR in both control and cKO mice (Fig. 7k-m), it caused dramatic increases in the OC.S/BS and OC.N/BPm in control mice, which were lost in vinculin cKO mice (Fig. 7n-r).

**Figure 7.**
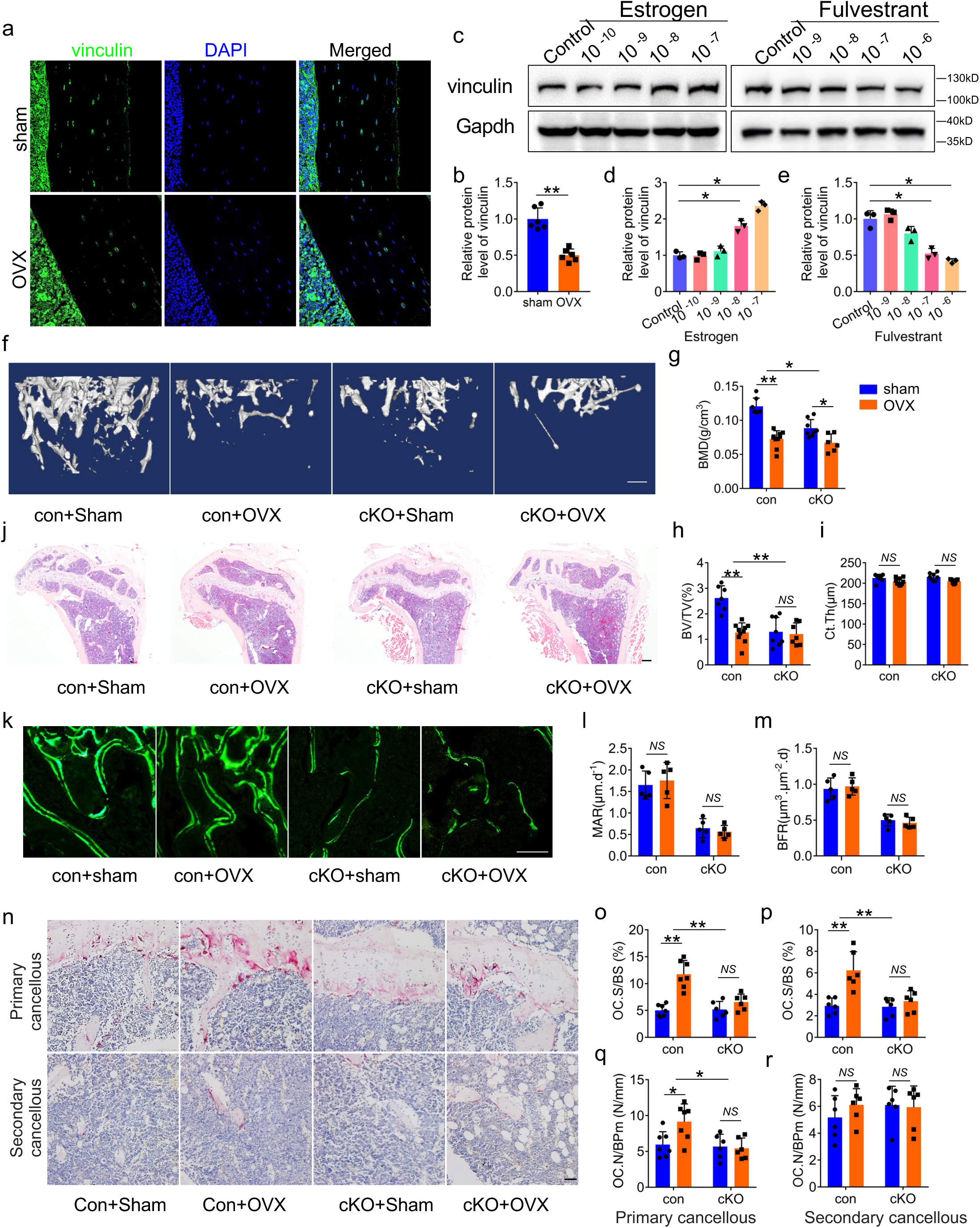
Estrogen controls vinculin expression in osteocytes and vinculin cKO mice are resistant to OVX-induced bone loss. **a, b** IF staining. Tibial sections of sham and OVX mice with an anti-vinculin antibody, Scale bars: 50 μm. Qauntitative data (b). *N* = 6 biologically independent replicates per group, *\*\*P* < 0.01, versus controls, two-way ANOVA. **c-d** Western blot analysis. MLO-Y4 cells were treated with increasing concentration of estrogen or fulvestrant (an estrogen receptor antagonist) for 24 h. Gapdh was used as a loading control. Quantitative data from three biologically independent replicates. Results were expressed as mean ± s.d., *\*P* < 0.05 versus control. **f** 3D reconstruction from μCT scans of femurs from 5-mo-old control and cKO female mice performed with sham or OVX surgeries. Scale bar, 250 μm. **g-i** Quantitative analyses of the BV/TV, Ct.Th and BMD. Results were expressed as mean ± s.d., *N* = 6 biologically independent replicates per group, *\*P* < 0.05, *\*\*P* < 0.01 versus controls, two-way ANOVA. **g** H/E staining of tibial sections. Scale bar, 200 μm. **k-m** Calcein double labeling. Representative images of 3-mo-old male femur sections (k). Sections of non-demineralized femurs were used for measurements of MAR and BFR for the metaphyseal trabecular bones. Scale bar, 50 μm. Results were expressed as mean ± s.d., *N* = 6 biologically independent replicates per group, *\*P* < 0.05 versus controls, two-way ANOVA. **n-r** TRAP staining. Tibial sections of 3-mo-male control and cKO mice were used for TRAP staining (n). Oc.S/BS (o, p) and Oc.N/BPm (q, r) of primary (o, q) and secondary (p, r) cancellous bones were measured using Image-Pro Plus 7.0. Scale bar, 50 μm. Results were expressed as mean ± s.d., *N* = 6 biologically independent replicates per group, *\*P* < 0.05, *\*\*P* < 0.01 versus controls, two-way ANOVA.

## Discussion

We summarize the main findings of the present study as follows. First, the expression of FA protein vinculin is drastically reduced in osteocytes in patients with age-related osteoporosis and in mice with estrogen deficiency. Second, deleting vinculin in Dmp1-expressing cells, i.e., primarily osteocytes and mature osteoblasts, causes a severe bone loss in both young and aged mice. Third, vinculin deletion impairs the bone mechanotransduction in mice. Fourth, vinculin expression in osteocytes is critically involved in estrogen control of bone mass in mice. Finally, vinculin regulates osteocyte adhesion and inhibits sclerostin expression to promote bone formation.

In this study, we demonstrate that vinculin increases bone mass and BMD by primarily promoting bone formation. Thus, vinculin loss in osteocytes and mature osteoblasts dramatically decreased the values of serum P1NP, BFR, MAR, OV/TV and mBV/TV, all osteoblastic parameters. These impairments lead to reduced bone formation and severe osteopenia. Of note, vinculin loss does not markedly alter osteoclast formation (OC.S/BS and OC.N/BPm) and bone resorption (serum CTX-1). This result on osteoclast formation is similar to that in mice lacking the other FA proteins pinch1/2 in Dmp1-expressing cells (Y. Wang et al., 2019), but in contrast to those in mice with deletion of FA protein kindlin-2 in the same cell type, which displayed increased osteoclast formation and bone resorption in part due to enhanced Rankl production in osteocytes (Cao et al., 2020). The reason(s) for this discrepancy are not clear.

Our findings from this study strongly suggest that vinculin inhibits sclerostin expression in osteocytes and mature osteoblasts to promote the osteoblast-mediated bone formation. Our in vitro and in vivo results demonstrate that vinculin-deficient osteocytes produce a large amount of sclerostin protein, which inhibits binding of Wnt ligands to the LRP5/6 receptors and, thereby, the Wnt/β-catenin signaling. The latter is a major player in control of the bone mass accrual in vertebrates. Consistently, vinculin loss reduces the level of active β-catenin protein and CFU-OB formation in bone marrows and bone formation in mice. Of particular significance, deleting *Sost* expression in Dmp1-expressing cells essentially reverses the low bone mass caused by vinculin loss. Thus, we provide important genetic evidence that FA protein vinculin plays a critical role in promoting bone formation by facilitating the Wnt/β-catenin signaling in osteoblastic cells.

At the molecular level, we provide intriguing evidence that vinculin loss promotes *Sost* expression through, at least in part, Mef2c, a major transcriptional regulator of the *Sost* gene (Collette et al., 2012; Kramer, Baertschi, Halleux, Keller, & Kneissel, 2012). Importantly, upregulation of sclerostin expression induced by vinculin loss is largely reversed by Mef2c KD in osteocytes. While vinculin loss does not alter the expression level of total Mef2c protein in osteocytes, it increases the Mef2c nuclear translocation and binding to the *Sost* enhancer *ECR5*. It is known that Mef2c binding to this enhancer is critical for *Sost* expression in osteocytes (Leupin et al., 2007). Our results suggest that binding of vinculin to Mef2c prevents the Mef2c nuclear translocation by retaining it in the cytoplasm.

While estrogen deficiency causes the postmenopausal osteoporosis, its mechanism(s) remain incompletely understood. We demonstrate that OVX mice express an extremely low level of vinculin protein in osteocytes in bone. Of particular significance, mice lacking vinculin in osteocytes are resistant to OVX-induced bone loss. We find that estrogen directly increases vinculin expression in osteocytes. Collectively, these results suggest that estrogen signaling favors bone formation by in part inducing vinculin expression in osteocytes. This knowledge improves our understanding of the molecular mechanism(s) underlying the postmenopausal osteoporosis.

Mechanical force is the most potent anabolic stimulus for bone formation. Osteocytes, the most abundant ed cells embedded in the mineralizing matrix in bone, are now believed to be the major sensor and mediator of mechanotransduction in bone (Qin et al., 2020). In this study, we demonstrate that vinculin expression in osteocytes is essential for skeletal response to mechanical loading to increase bone formation. It should be noted that loss of other FA proteins Kindlin-2 or pinch in osteocytes caused a similar defect in the anabolic effects of mechanical loading on bone in mice (Qin, Fu, et al., 2021; Y. Wang et al., 2019). Collectively, these results suggest a critical role of the FA signaling pathway in mediation of the mechanotransduction in bone. Notably, previous studies demonstrate that mechanical force promotes vinculin activation through conformational changes and a loss of tension causes vinculin to be rapidly inactivated (Carisey et al., 2013).

Finally, we propose a working model regarding how FA protein vinculin expression in osteocytes embedded in the bone matrix controls bone formation (Fig. 8). In young and adult animals, vinculin is highly expressed in osteocytes where it, probably together other FA proteins, such as talin and actin, forms a complex with Mef2c in the cytoplasm. Complex formation prevents the nuclear translocation of Mef2c, leading to reduced expression and production of sclerostin and, thereby, increased bone formation. In contrast, in the presence of vinculin deficiency in osteocytes, which is related to aging and estrogen deficiency, the nuclear translocation of Mef2c and its binding to the *Sost* enhancer *ECR5* are enhanced, thus stimulating Sost expression and inhibiting bone formation. In summary, in this study, we demonstrate that vinculin plays an important role in control of bone formation and may be a useful target for the prevention and treatment of aging and estrogen deficiency induced osteoporosis.

**Figure 8.**
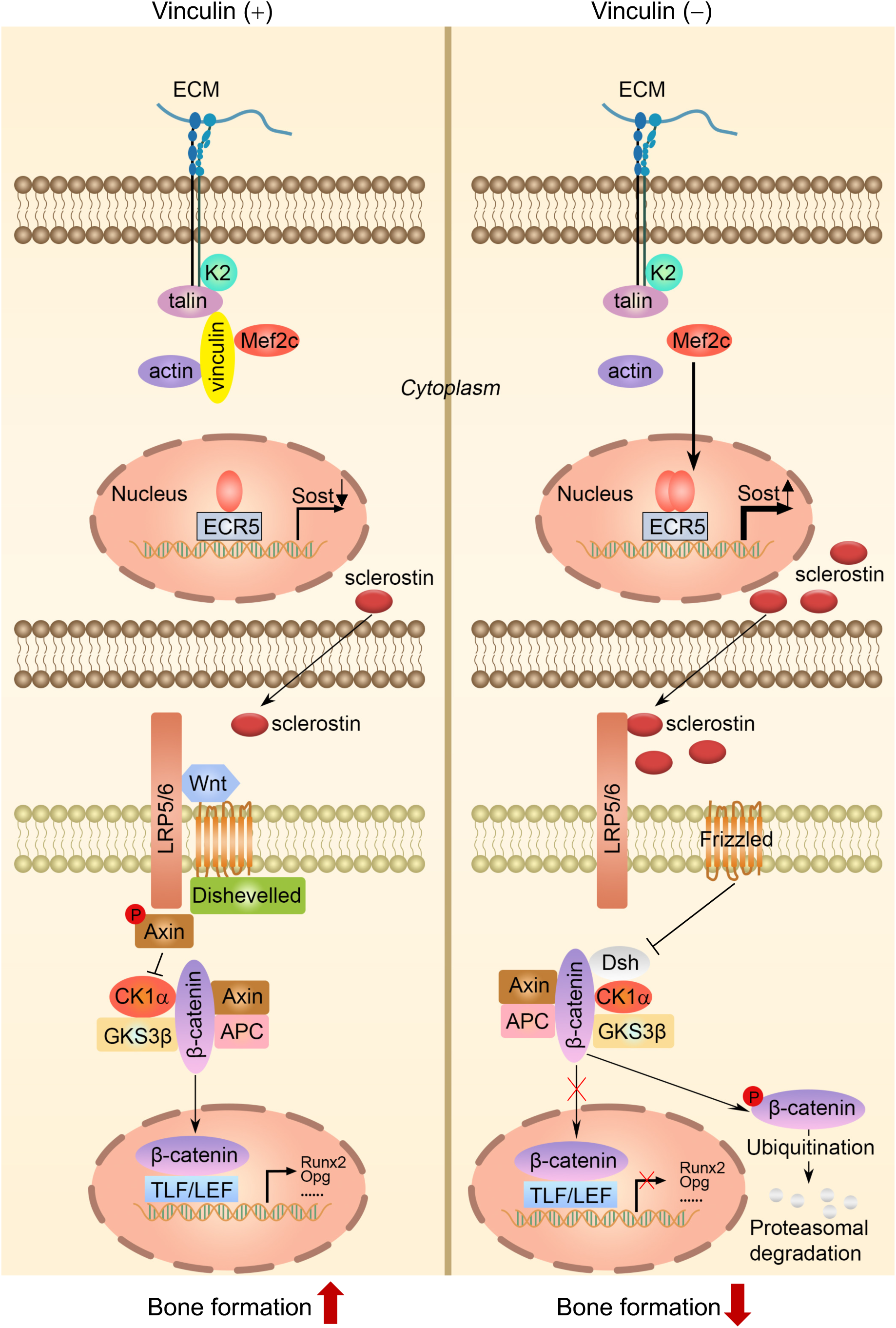

## MATERIALS AND METHODS

### Animal studies

*Dmp1-Cre* mice were generously provided by Dr. Jian Q Feng (Baylor College of Dentistry) (J. Q. Feng et al., 2003). *Vcl ^fl/fl^* mice were purchased from the Jackson Laboratory. The experimental animals used current studies were obtained using a 2-step breeding strategy. Homozygous *Vcl ^fl/fl^* mice were first crossed with *Dmp1-Cre* transgenic mice to generate *Dmp1-Cre*; *Vcl ^fl/+^* mice. These offspring were then crossed with *Vcl ^fl/fl^*to generate *Dmp1-Cre*; *Vcl ^fl/fl^* mice. The generation of the *Dmp1-Cre*; *Vcl ^fl/fl^*; *Sost ^fl/fl^*mice was as follow: *Dmp1-Cre*; *Vcl ^fl/fl^* mice were first crossed with *Sost^fl/fl^* mice to generate *Dmp1-Cre*; *Vcl ^fl/+^*; *Sost ^fl/+^* mice. These offspring were then intercrossed with *Vcl ^fl/fl^*; *Sost ^fl/fl^* mice to generate the *Dmp1-Cre*; *Vcl ^fl/fl^*; *Sost ^fl/fl^* mice and other genotypes. In vivo tibial loading experiments were performed as previously described (Qin, Fu, et al., 2021). All animal experiments in this study were approved by the Institutional Animal Care and Use Committees (IACUC) of the Southern University of Science and Technology.

### Tissue samples

Human cancellous tissues and their corresponding adjacent normal controls were collected from the Shenzhen Hospital of Guangzhou University of Chinese Medicine. Written informed consent was obtained from all patients. The study has been approved by the Shenzhen Hospital of Guangzhou University of Chinese Medicine, China (NO. GZYLL(KY)-2022-027). Specimens were collected and stored in formaldehyde after surgery.

### Micro-computerized tomography (μCT) analysis

Fixed non-demineralized femur, tibia, spine or skull were used for μCT analysis in the Experimental Animal Center of Southern University of Science and Technology using a Bruker μCT (SkyScan 1176 Micro-CT, Bruker MicroCT, Kontich, Belgium) following the standards of techniques and terminology recommended by the American Society for Bone and Mineral Research (ASBMR) (Bouxsein et al., 2010).

### Histological evaluation and bone histomorphometry

For histology, bone tissues were fixed in 4% paraformaldehyde (PFA) at 4°C for 24 h, decalcified in 10% EDTA (pH 7.4) for 21 d, and embedded in paraffin after dehydration. Bone sections were used for hematoxylin and eosin (H/E) and tartrate-resistant acid phosphatase (TRAP) staining using our standard protocols (Y. Wang et al., 2019; C. Wu et al., 2015).

### Immunohistochemistry (IHC) and immunofluorescence (IF) staining

For IHC staining, 5-μm sections were stained with antibodies or control IgG using the EnVision+System-HRP (DAB) kit (Dako North America Inc, Carpinteria, CA, USA) as previously described (Cao et al., 2020). IF staining of bone sections and MLO-Y4 cells was performed as we previously described (Zhu et al., 2020).

### BMSC culture

Primary bone marrow stromal cells (BMSC) were isolated from femurs of 3-month-old male mice as previously described (Yan, Gao, Yao, Ling, & Xiao, 2022). Briefly, bone marrow cells were cultured on 60-mm cell culture dishes in α-MEM (Hyclone, USA) containing 15% FBS. After 48 h, the non-adherent cells were removed. On the seventh day, the cells trypsinized for subsequent experiments. Colony forming unit-fibroblast (CFU-F) assay and colony forming unit-osteoblast (CFU-OB) assay were performed as we previously described (Lei et al., 2020).

### ELISA assays

Serum levels of P1NP were measured using the RatLaps EIA Kit (cat# AC-33F1) according to the manufacturer’s instruction (Immunodiagnostic Systems Limited). Serum levels of CTX, degradation products from type I collagen during osteoclastic bone resorption, were measured using the RatLaps EIA Kit (cat# AC-06F1) according to the manufacturer’s instruction (Immunodiagnostic Systems Limited).

### RNA extraction and qRT-PCR analysis

RNA isolation and quantitative real-time RT-PCR analysis were performed as we previously described (H. Gao et al., 2019). The specific primers for gene expression analysis were listed in Supplementary Table 1.

### Western blot analysis

Western blot analysis was performed as previously described (Y. Wang et al., 2021). Briefly, whole-cell lysates were prepared in RIPA lysis buffer (Sigma, USA) and aliquots of 15 μg protein were separated by SDS-PAGE and blotted onto a polyvinylidene fluoride (PVDF) membrane (Millipore, MA, USA). Membranes were blocked at room temperature for 15 min in QuickBlock™ Western (Beyotime), followed by an overnight incubation at 4 °C with specific antibodies. After incubation with appropriate HRP-conjugated secondary antibodies (ZSGB-bio), blots were developed using an enhanced chemiluminescence (ECL Kit, BIORAD) and exposed in ChemiDoc XRS chemiluminescence imaging system. Antibodies used in this study are listed in Supplementary Table 2

### Co-immunoprecipitation (Co-IP) assay

Co-IP assay was performed as previously described (Fu et al., 2020). Briefly, cells were incubated for 10 min at 4 °C in RIPA buffer (Sigma, USA). After a centrifugation at 12,000 × g for 10 min at 4 °C, the supernatant was first incubated with corresponding primary antibody overnight and then with Protein A/G Magnetic Beads at room temperature for 1 h. DynaMag™-2 Magnet (Thermo Fisher) was used to collect dynabeads-antigen-antibody complex. The complex was washed with IP buffer three times and resuspended with 60 μl 1× loading buffer and cooked at 95 °C for 5 min, followed by SDS-PAGE and western blotting.

### Chromatin immunoprecipitation (ChIP) assay

ChIP assay was performed using the ChIP Kit (Abcam, ab500) as previously described (X. Wang et al., 2020). Briefly, 1 × 10^7^ cells were collected in lysis buffer and sonicated to generate chromatin samples with average fragment sizes of 100–500 bp. Cell lysates were incubated with the indicated antibodies or IgG overnight at 4°C. Then, the supernatants were mixed with the blocked Protein A/D sepharose beads to collect the antibody–chromatin complexes. After washing 4 times, the immunoprecipitated DNA was eluted and purified for the subsequent qPCR analysis.

### Statistical Analyses

The sample sizes for all experiments conducted in this study were determined based on our previous experience on similar studies. Mice used in this study were randomly grouped. The IF, IHC and histology were conducted and analyzed under double-blind conditions. Statistical analyses were completed using the Prism GraphPad. Two-way ANOVA and unpaired two-tailed Student’s *t*-test was used as indicated. *P* < 0.05 was considered statistically significant.

### Data availability

All data generated or analysed during this study are included in the manuscript and supporting file; Source Data files have been provided for Figures.

## Acknowledgments

The authors acknowledge the assistance of Core Research Facilities of Southern University of Science and Technology. This work was supported, in part, by the National Key Research and Development Program of China Grant (2019YFA0906004), the National Natural Science Foundation of China Grants (82230081, 82250710175, 81991513, 81630066, 81870532, 82004395), the China Postdoctoral Science Foundation (2021M691435) and the Guangdong Provincial Science and Technology Innovation Council Grant (2017B030301018).

## Author contributions

Study design: GX and YW. Conducting study, data collection and analyses: YW, JH, SL, LQ, DH, PZ and SH. Data interpretation: GX, YW, XZ and DC. Drafting the manuscript: GX and YW. YW and GX take the responsibility for the integrity of the data analysis.

---

**Supplementary Table 1:**
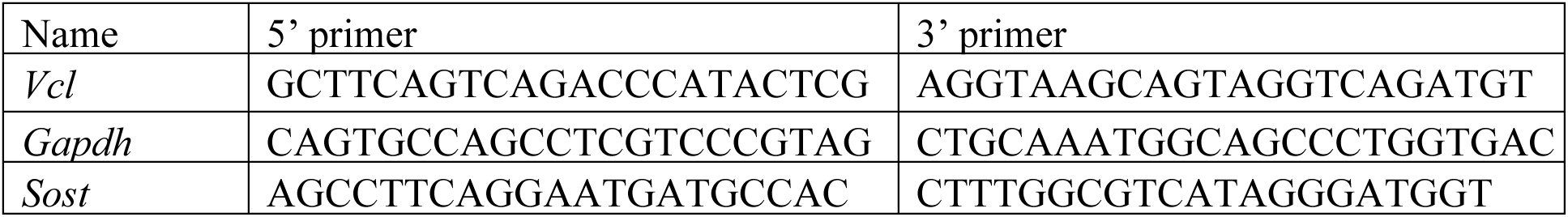
Mouse real-time RT-PCR (qPCR) primers.

**Supplementary Table 2:**
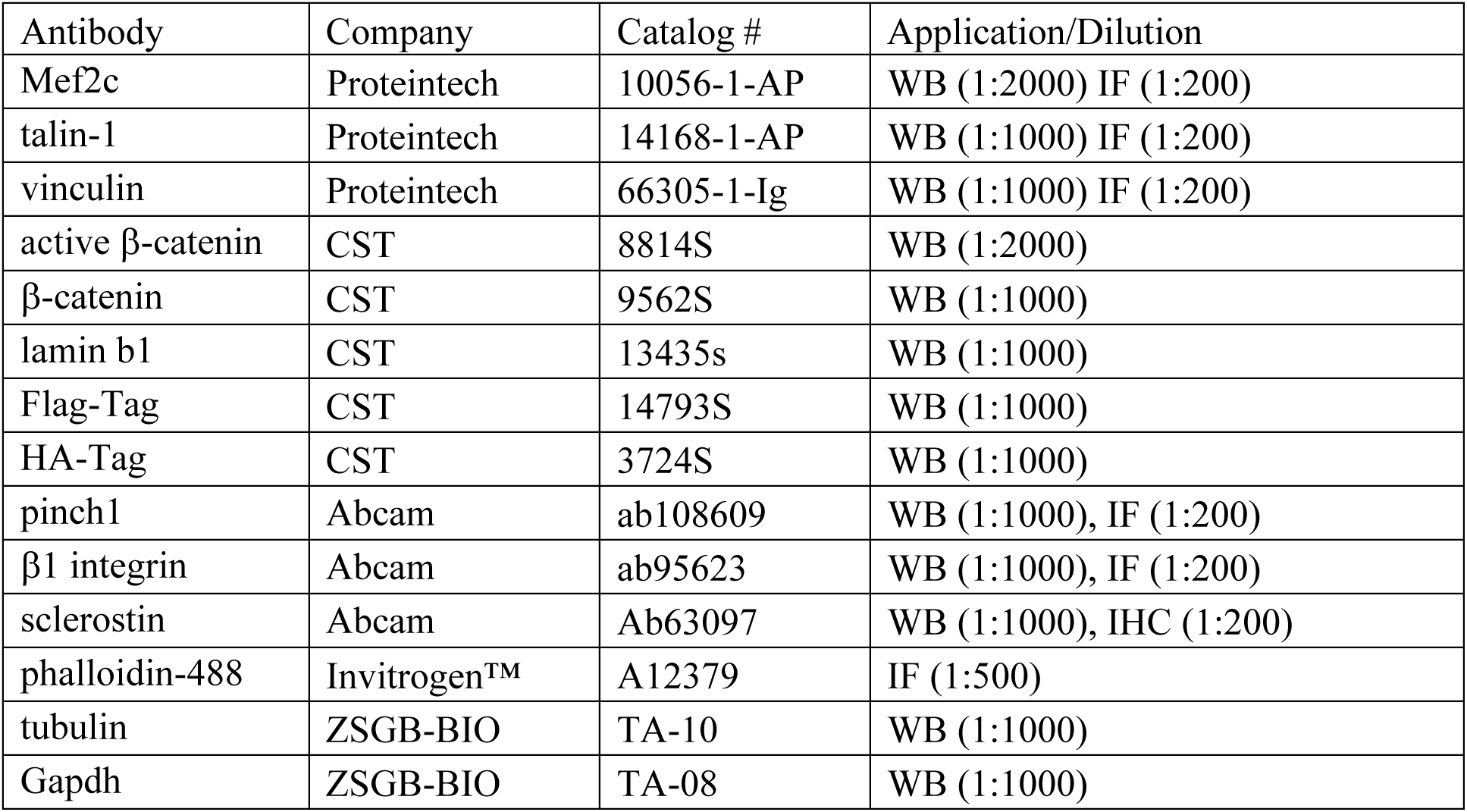
Antibody information.

